# Anisotropic Interaction and Motion States of Locusts in a Hopper Band

**DOI:** 10.1101/2021.10.29.466390

**Authors:** Jasper Weinburd, Jacob Landsberg, Anna Kravtsova, Shanni Lam, Tarush Sharma, Stephen J Simpson, Gregory A Sword, Jerome Buhl

## Abstract

Swarming locusts present a quintessential example of animal collective motion. Juvenile locusts march and hop across the ground in coordinated groups called hopper bands. Composed of up to millions of insects, hopper bands exhibit aligned motion and various collective structures. These groups are well-documented in the field, but the individual insects themselves are typically studied in much smaller groups in laboratory experiments. We present the first trajectory data that detail the movement of individual locusts within a hopper band in a natural setting. Using automated video tracking, we derive our data from footage of four distinct hopper bands of the Australian plague locust, *Chortoicetes terminifera*. We reconstruct nearly twenty-thousand individual trajectories composed of over 3.3 million locust positions. We classify these data into three motion states: stationary, walking, and hopping. Distributions of relative neighbor positions reveal anisotropies that depend on motion state. Stationary locusts have high-density areas distributed around them apparently at random. Walking locusts have a low-density area in front of them. Hopping locusts have low-density areas in front and behind them. Our results suggest novel insect interactions, namely that locusts change their motion to avoid colliding with neighbors in front of them.

## 1 Introduction

Locust hopper bands exhibit a striking example of collective motion in insects. Without a complex social structure, juvenile locusts self-organize into a range of patterns from columnar streams to planar fronts that appear to serve ecological functions for the group, such as migration and foraging [1, 2]. Similar collective motion is observed in schools of fish [3], flocks of birds [4, 5], and even herds of ungulates [6]. These natural phenomena have produced a field of research centered on the idea that a group can attain collective goals without centralized instruction.

Most modern models of collective motion rely on a few rules for simple interactions between individuals, and so are fundamentally similar to their predecessors [7, 8, 9]. These interactions are often characterized as attraction, repulsion, and alignment of the direction of motion. Empirical studies by biologists, ecologists, and physicists have provided evidence supporting these models and informing these interaction rules for specific species, for instance in starlings [10] and golden shiners (fish) [11]. Buhl et al. [12] has previously made observations for locust interactions, which we aim to build on here.

Locusts and their motion are the subject of studies ranging from empirical work [13, 14, 15, 16, 1, 12] to theoretical modeling [17, 18, 19]; for a review see the work of Ariel and Ayali [20]. Locust behavior and motion is grounded in their biology. Locusts exhibit phase polyphenism, a phenomenon whereby an individual can exhibit two distinct phases of behavior (and for some species also distinct morphologies). In the solitarious phase, locusts typically avoid each other and forage individually. Crowding by conspecifics triggers a transition to the gregarious phase in which individuals gather in social aggregations [21]. When composed of juveniles, these aggregations are called *hopper bands* because locust nymphs, whose wings have not fully developed, hop and walk across the ground. Hopper bands are often composed of hundreds of thousands of locusts all moving as a collective [16, 1]. Researchers study how the persistence, motion, and shape of a hopper band develop from the interactions between individuals through the lens of collective motion.

Relatively simple collective motion models have successfully reproduced collective patterns of hopper bands observed in the field. Dkhili et al. [17] demonstrated that both columns and fronts can be achieved by varying individual-level parameters in a model that incorporated only local interactions between locusts. A similar approach was taken by Bach [18], with a realistic number of individuals and parallel computing approach. An alternative approach by Bernoff et al. [19] incorporated individual locust interactions with food resources and showed that dense fronts are typical when sufficient food is present. Modelers often rely on hypothesized interactions at the individual level. The most common assumption is the simplest; that individual interactions are isotropic, that is, interactions depend only on the distance between individuals and not on their relative positions. One notable exception is the escape-and-pursuit model [22], which hypothesizes that proximate behavioral responses are the result of ultimate selection pressures related to cannibalism risk [14] (chasing those in front and fleeing from those behind). In this modeling framework, locusts are likely to have neighbors directly in front or behind them [23], but this has not been observed empirically.

A majority of the empirical work studying the motion of hoppers is conducted through either laboratory experiments focused on individuals [13, 14, 15] or field observations of the group as a whole [16, 1, 12]. These field observations date back to Clark [16] and Ellis and Ashall [24] and are mainly qualitative in nature, for instance noting the shape of the hopper band at different times of day or in varying vegetation cover. Many laboratory studies are conducted by placing a relatively small number of locusts (less than a hundred) in an arena and observing their motion. While these empirical studies have advanced our understanding of the mechanics of locust motion and interaction, there is still a particular dearth of data on individual interactions during collective motion within a group of a naturally occurring size.

We present and analyze the first trajectory data of individual locusts moving within a hopper band. We study four hopper bands of the Australian plague locust, *Chortoicetes terminifera*, from an outbreak in 2010 near Hillston, New South Wales, AUS. We recorded video of locusts moving across the ground using cameras mounted on tripods (see Sample video footage for a sample). Using automatic tracking software (TrackMate [25]) we extracted 19 687 individual trajectories by linking 3 369 723 locust positions. Our analysis of these trajectories suggests that locusts adjust their motion to avoid neighbors ahead of them, providing evidence of a novel locust-locust interaction for collision avoidance. These results add to the understanding of individual interactions in marching locusts and provide valuable insight for modelers seeking to reveal the mechanisms behind the collective motion of the swarm.

## 2 Results

Our data set was extracted from recordings of four bands of Australian plague locust, *Chortoicetes terminifera*. The data consists of 3 369 723 locust positions linked into 19 687 trajectories from 24 300 frames (twenty-seven minutes) of video. Three sample trajectories are shown in Figure 1 (top), with a still image and processed data from Appendix A. From all trajectories we inferred a total of 3 332 137 heading directions and individual speeds; Figure 1 (left) shows a histogram of these speeds. We constructed a plot of the relative density around a focal individual from 19 407 719 nearby neighbor positions, shown in Figure 1 (right). The anisotropy apparent in this plot contrasts with previous findings by Buhl et al. [23] and motivates deeper investigation. To this end, we classified the data into distinct motion states (stationary, walking, and hopping) using statistical learning. We partitioned the relative neighbor data by motion state of the focal individual to reveal differences in anisotropy depending on motion. Of note, in the remainder of the paper, we often use “locust” to refer to the Australian plague locust (APL) specifically and acknowledge that some results may be species specific.

**Figure 1:**
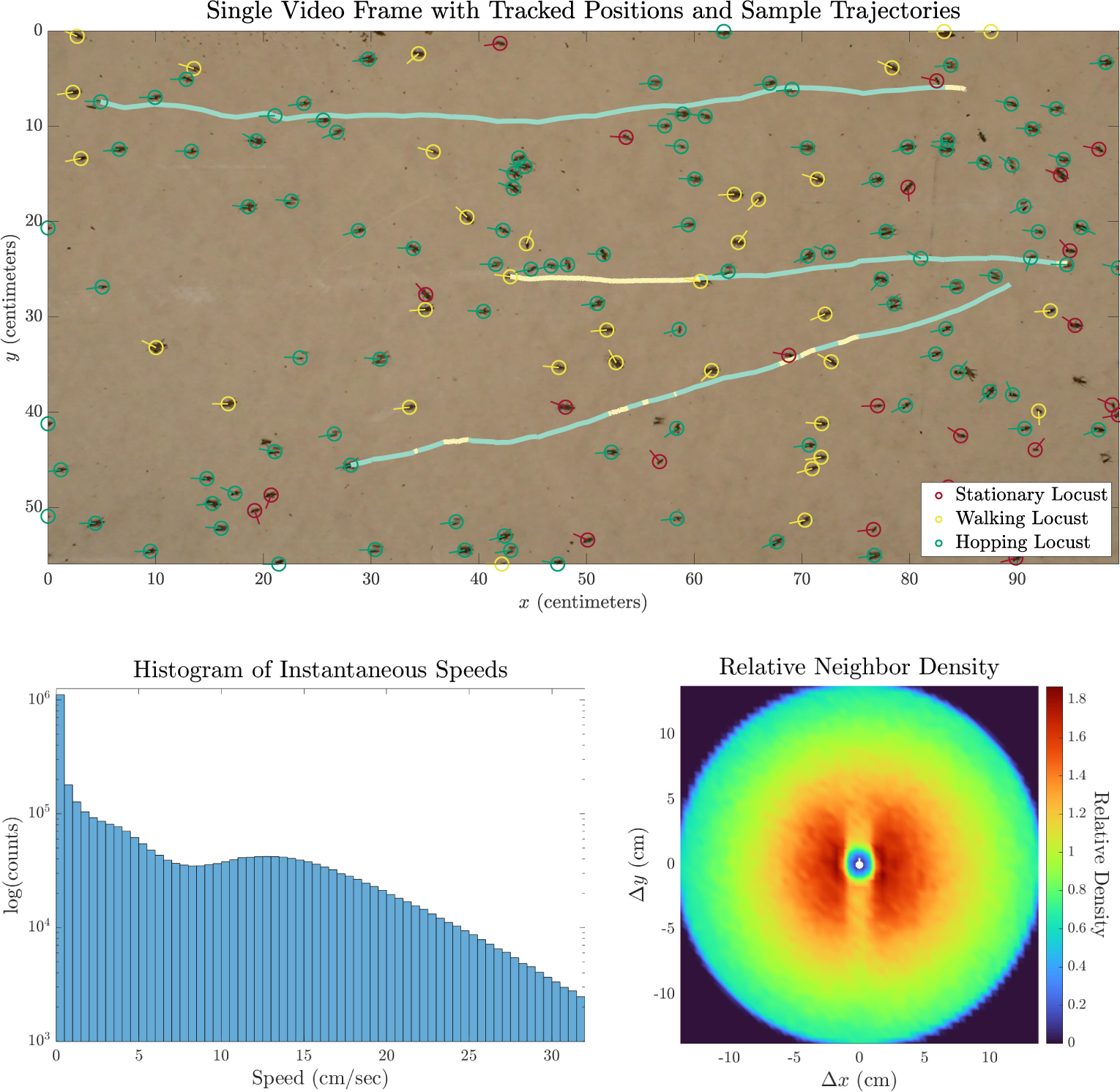
Single frame of video with processed data (top), histogram of speeds (left), and plot of relative neighbor density (right). The image (top) is taken from the video with processed data in Appendix A and augmented showing the trajectories of three sample locusts with color denoting the motion state. The distribution of speeds (left) is bimodal, with peaks near 0 and 13 cm s^−1^. From approximately 2–5 cm s^−1^, the counts decrease exponentially (linear decrease on the logarithmic scale). The relative neighbor density (right) is computed from 19 407 719 relative neighbor positions around a representative focal locust (white marker) positioned in the center and oriented facing upwards, along the vertical axis. Density is highest (red) within a radius of 5 cm and lowest (blue) in a central disc with radius approximately 1 cm. The density is not rotationally symmetric; at distances of 1–7 cm there is a noticeable decrease in density (orange) directly in front (above) and behind (below) the focal locust. At distances greater than 7 cm, the density appears to be rotationally symmetric.

### 2.1 Collective marching

All four bands exhibited collective marching behavior. The mean density across all recordings 141.6 locusts*/*m^2^ is well above the established 20 locusts*/*m^2^ threshold for marching [13]. Measurements of group alignment (polarization and entropy index) agree with previous measurements of marching locusts in the field [12]. Polarization is the length of the average of the direction vector and our entropy index is an adaptation of Boltzmann entropy, see Measuring collective marching and Appendix F for details. We compute a mean polarization of 0.82 and mean entropy index of 0.76 across all bands.

We present values for each band in Appendix B (Table 2) and direct measurements for a regular subsample of frames (every fifth frame) as scatter plots in Figure 2. The density of each band is relatively well-cluster, as expected. There is a distinct decrease in entropy with increasing density and a high variance in polarization for densities below 200 locusts*/*m^2^. See [12] for an in-depth analysis of similar trends.

**Figure 2:**
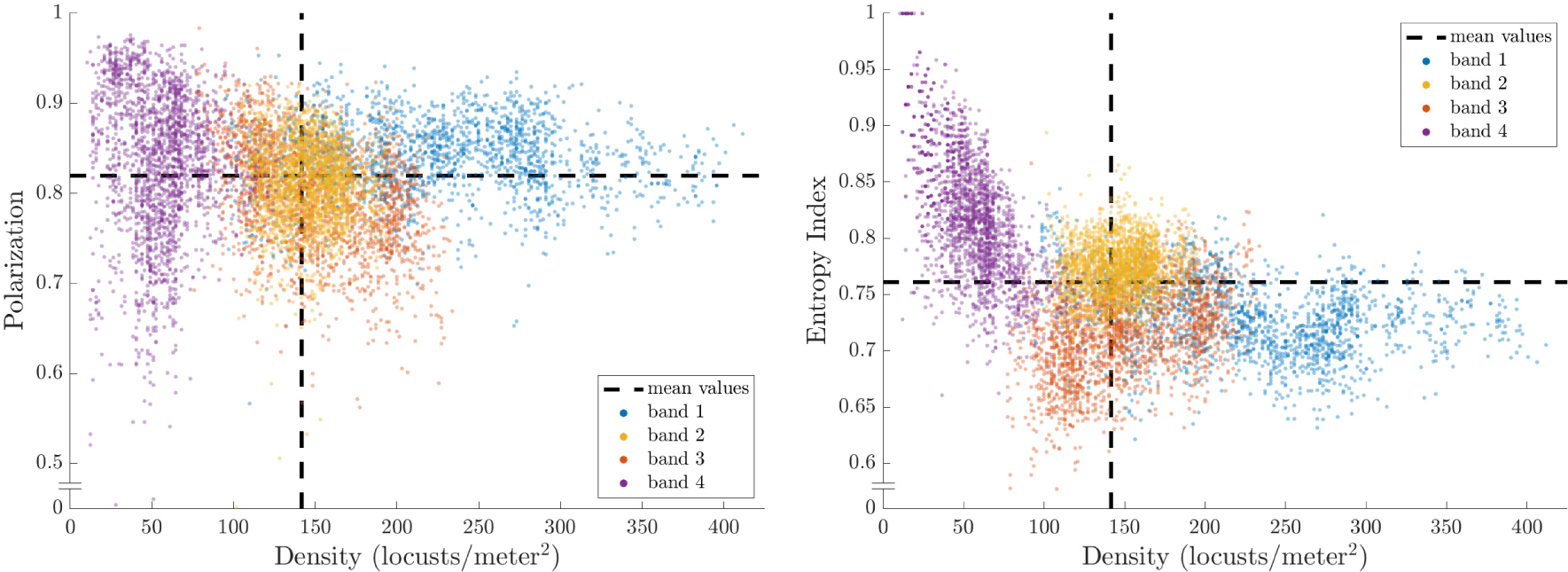
Measures of collective alignment plotted against density for all bands (distinguished by color). Polarization (left) is highly variable for densities less than 200 locusts*/*m^2^. Entropy (right) shows a marked decrease with increasing density, consistent with [12]. Data plotted is a regular subsample of frames (every fifth frame), with mean values of the full data set (dashed lines). Essentially all measurements lie within ranges associated with collective marching for hopper bands.

### 2.2 Individual speed and motion state

Of the 3 369 723 locust positions, we computed 3 332 137 individual speeds, presented in Figure 1 (left). (The discrepancy in the number of positions and speeds is due to the start and end of trajectories.) Each speed was classified into one of three motion states by a support vector machine. Our classification uses four summary statistics computed in a moving time window; further details are provided in Classifying motion state. These motion states divide the data into 26.0 % stationary, 24.7 % walking, and 49.2 % hopping. While there is some variation between bands in the composition by motion state, this variation does not appear to be correlated with density or group alignment. We computed mean speeds for each motion state and found (0.2 ± 0.7) cm s^−1^ (stationary), (2.7 ± 2.2) cm s^−1^ (walking), and (11.8 ± 9.3) cm s^−1^ (hopping). Each mean speed has plus/minus one standard deviation. Motion state and speed data are presented for each band in Appendix B (Table 3). See Sample video footage for a visualization showing results of the motion state classification.

Figure 3 shows histogram plots for the speed distributions divided by motion state. Note the logarithmic scale on the vertical axis. Speeds of stationary locusts essentially all fall into the first two bins 0–1 cm/sec. Higher speeds make up less than 2.5% of all data classified as stationary. The number of walking speeds have no such dramatic peak near 0, instead decreasing slowly until 5 cm s^−1^, then decreasing super-exponentially (concave down on the logarithmic plot). Speeds of hopping locusts are by far the most widely distributed with two peaks at 0 and 13 cm s^−1^, matching our observation that hopping locusts often pause briefly between two jumps. A minimum around 5 cm s^−1^ separates these two maxima and the hopping speeds decrease super-exponentially after the second. Comparing these three plots to Figure 1 (left), we observe that our motion state assignment has cleanly divided the data into three distributions with unique features that were each visible in the full distribution.

**Figure 3:**
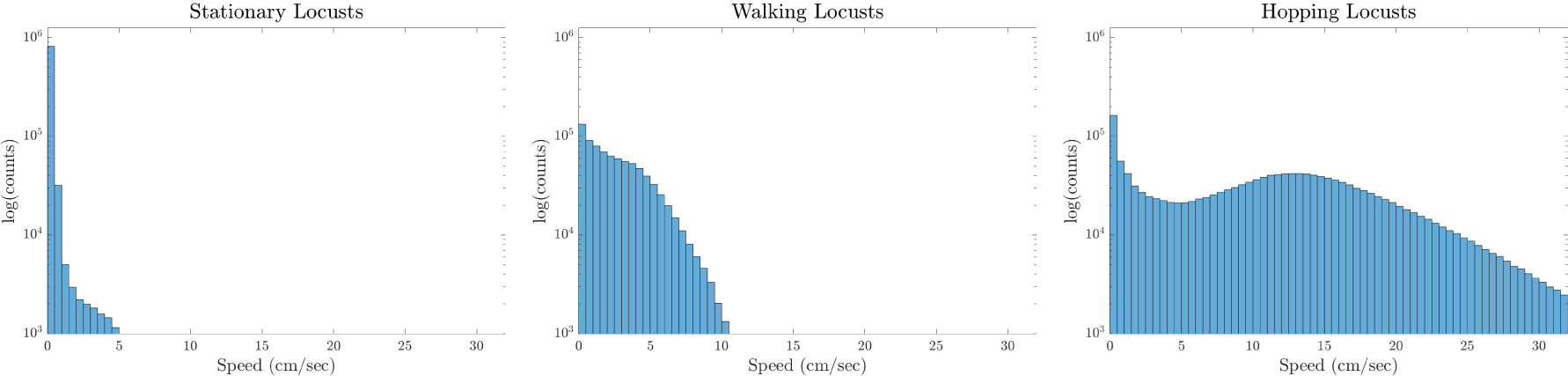
Histograms of speeds for stationary (left), walking (center), and hopping (right) locusts. All appear with the same log-scaled vertical axis. Matching intuition, the speeds of stationary locusts are tightly grouped near 0 cm s^−1^ (speeds above 1 cm s^−1^ make up less than 2.5% of all stationary data). Walking locusts have speeds between 0 and 10 cm s^−1^ with a steeper decline in numbers after 5 cm s^−1^. Hopping locusts have a bimodal distribution with one peak near 0 and the other near 13 cm s^−1^, as we expect from the pattern of pausing between hops observed in our recordings.

### 2.3 Anisotropy in relative neighbor density

Figure 1 (right) demonstrates that the relative neighbor density is not isotropic, i.e. not rotationally symmetric, particularly at distances of 1–7 cm from the focal individual. At distances greater than 7 cm, the density appears isotropic. We focus next on a smaller square around the focal locust so as to exclude the isotropic region at larger radii.

In Figure 4, we plot the relative neighbor densities around focal locusts that are stationary, walking, and hopping. In each plot, the focal locust is positioned in the center (Δ*x* = Δ*y* = 0) and faces upwards (along the Δ*y*-axis). The highest relative neighbor densities are indicated by red, intermediate densities are shown in green, and the lowest neighbor densities appear in blue. For all three motion states, there is a roughly circular area of low density around the focal individual with a radius of approximately 1 cm. Past a radius of 7 cm, the plots are roughly isotropic (rotationally symmetric) and similar between motion states.

**Figure 4:**
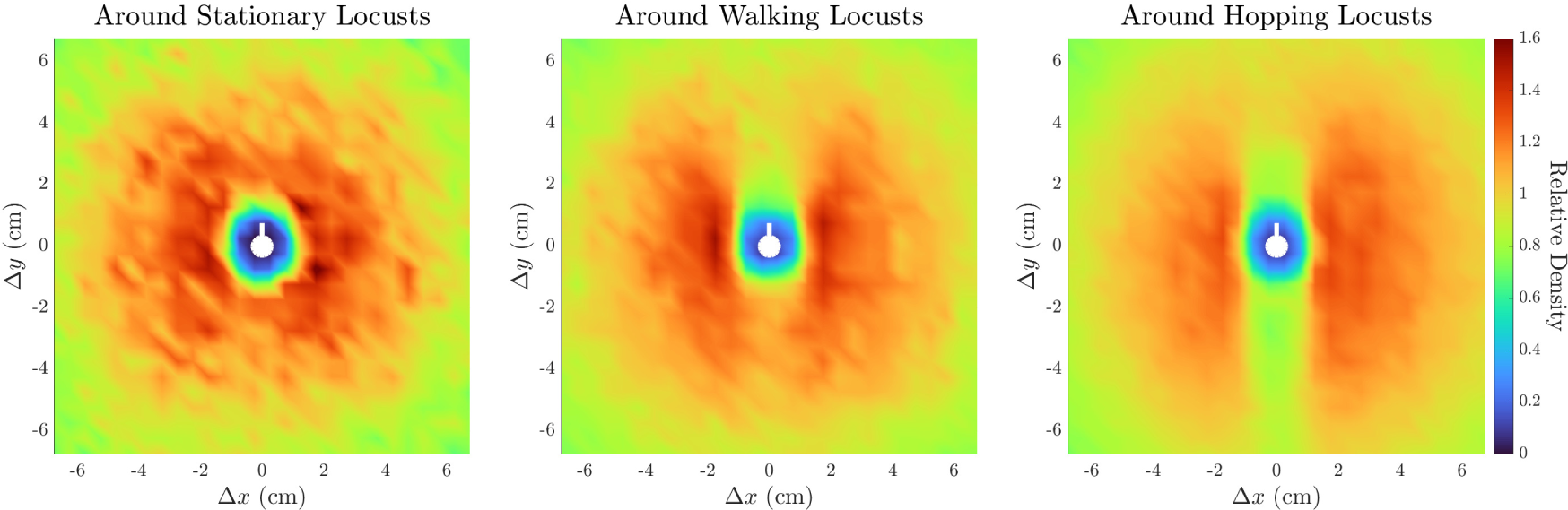
Relative neighbor density around locusts that are stationary (left), walking (center), and hopping (right). A representative focal locust (white marker) is positioned in the center of each plot and oriented facing upwards along the vertical axis. Areas of highest density (red) are distributed apparently at random around a stationary focal locust but highly anisotropic around walking and hopping focal locusts. A notable sector-shaped area of lower density (green-orange) lies immediately in front (above) the walking focal locust. A longer and narrower strip-shaped area of low density (green) lies in front and behind the hopping focal locust.

The visual differences in relative density plots between motion states in Figure 4 are striking. For stationary focal locusts, the relative neighbor density is mostly isotropic, with localized spots of high density (red) distributed apparently at random angles around the focal locust. For walking locusts, there is a distinct area of lower density ahead of the focal individual. This void has the approximate shape of a 45^◦^-sector. The highest relative neighbor densities are directly to the left and right of the focal individual at a distance just less than 2 cm. For hopping locusts, there is a strip-shaped area of low density ahead and behind the focal individual. This void appears to divide an otherwise circular area of high density that decreases with distance from the center. The high-density area to the right of the focal individual is larger than on the left, which we attribute to a density gradient in recordings of bands 1 and 3. The observed low-density sector in front of walking locusts and the strip both in front and behind hopping locusts are novel anisotropies for neighbor densities in the APL and other locust species.

We quantified these anisotropies by examining the angular distribution of neighbors within 7 cm. First, the Hodges-Ajne test for uniformity confirms that, for a focal locust in any motion state, the distribution of neighbor angles is not drawn from the uniform distribution of angles; we compute *p*-values less than 10^−10^ (stationary), less than 10^−128^ (walking), and less than machine error (hopping). We characterize the degree of this nonuniformity by computing trigonometric moments 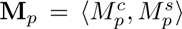, which we record in Appendix B (Table 4). Overall anisotropy |**M**_p_| is significantly larger for the distributions around walking and hopping locusts (|**M**_1_| = 0.0173, 0.0203 and |**M**_2_| = 0.0248, 0.0358) than around stationary locusts (|**M**_1_| = 0.0040 and |**M**_2_| = 0.0146). This quantitatively confirms what Figure 4 shows visually – that the distributions around moving locusts are less isotropic than around stationary locusts.

In Figure 4 we noted two forms of anisotropy around moving locusts. First, the sector of lower density in front of walking locusts is unimodal and therefore measured by **M**_1_. We compute the *front-back asymmetry* as 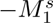 = **M**_1_ · (0, −1) to measure its size along the apparent axis of asymmetry (vertical). The negative captures the lower density in front and higher density behind. We point out the extreme disparity in 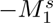 between the distributions around walking (0.0173) and stationary (0.0023) focal locusts. Second, both walking and hopping locusts have areas of high density on either side. This is measured by **M**_2_ and we compute the *four-fold anisotropy* 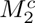 = **M**_2_ · (1, 0) to measure its size along the apparent axis of high density (horizontal). We highlight the extreme disparity in *M^c^* between the distributions around hopping (0.0357) and stationary (0.0142) focal locusts. These values are reported in bold in Appendix B (Table 4), along with other detailed anisotropy quantities.

Finally, in Figure 5 we examined how front-back asymmetry 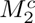 (dashed blue) and four-fold anisotropy 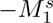 (solid orange) depend on distance from the focal individual. We computed 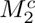 and 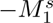 for subsets of neighbors from overlapping annuli of width Δ*r* = 1 cm at intervals of *r* = 0.25 cm. In each annulus, we normalized the anisotropy by scaling with the ratio of the density in that annulus to the density in the complete disc with radius 14 cm. For a measure of distance from uniformity, we plotted the same quantities computed analytically for a uniform distribution. The uniform distribution of angles on the circle has 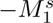 = 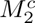 = 0 (black) and standard deviation 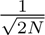, where *N* is the number of neighbor positions in the current annulus. Gray shading represents ±5 standard deviations.

**Figure 5:**
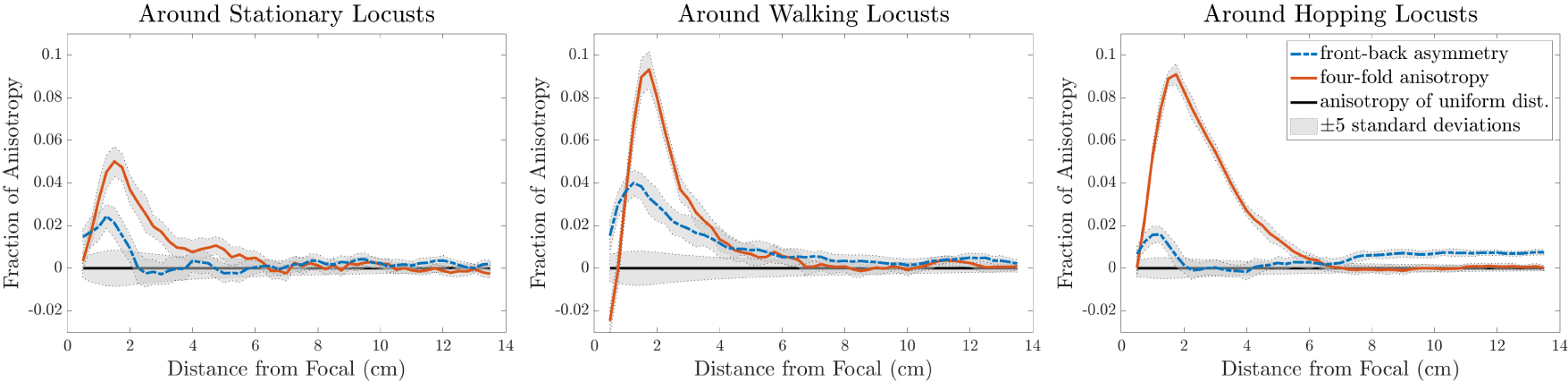
Anisotropy around stationary (left), walking (center), and hopping (right) locusts as functions of distance from the representative focal individual. Each plot shows front-back asymmetry 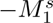 (dashed blue), four-fold anisotropy 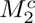 (solid orange), and anisotropy computed analytically for a uniform distribution 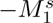 = 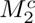 = 0 (black). Gray shading represents ±5 standard deviations. Both measures of anisotropy around stationary locusts are approximately half of the same measure around moving locusts and decrease quickly towards uniformity. Front-back asymmetry (dashed blue) peaks before decreasing slowly around walking locusts, while it is generally smaller around hopping locusts. Four-fold anisotropy (solid orange) around both walking locusts and hopping locusts peaks at a radius of 2 cm, then decreases quickly around walking locusts and slowly around hopping locusts. At distances above 8 cm, there is small but significant front-back asymmetry around hopping locusts.

For a given distance, both measures of anisotropy around stationary locusts (left) are small – approximately half – compared to the same around moving locusts (center, right). Front-back asymmetry 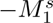 (dashed blue), is more than twice as large around walking locusts (center) than around either other motion state. Smaller front-back asymmetry around hopping locusts at short distances may be attributed to their high movement speeds. Specifically, any neighbors or open spaces ahead of them will be behind them a fraction of a second later; our time-aggregated plots manifest this as front-back symmetry around hopping locusts. Four-fold anisotropy 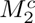 (solid orange) peaks around both walking locusts and hopping locusts (right) at a radius of approximately 2 cm, but decreases quickly around walking locusts and more slowly around hopping locusts. Both anisotropy quantities decrease to near 0 at a distance between 6–8 cm for all motion states. Around hopping locusts, we notice a small increase in front-back asymmetry for distances greater than 8 cm. Values for distances smaller than 0.5 cm are computed from relatively few neighbors, resulting in the initial negative value of *M^c^* around walking locusts. These computations quantify our visual observations of Figure 4. Moreover, they reveal lengthscales where the anisotropies are strongest.

## 3 Discussion

Insight into animal collective behavior develops from the feedback between theoretical studies of universal models and empirical studies of the animals themselves. The modeling approach seeks to reproduce a range of incredible structures, patterns, and group strategies based on simple and often local interactions between individuals. Meanwhile, the empirical work aims to uncover the specifics of individual-level behaviors. One challenge of conducting empirical studies has been to capture individual behavior in the midst of large and often dense groups of animals. This has been a particular difficulty for locusts due to the rarity of their swarming, small size, and disorder compared to larger animals. Identifying coherent signal amidst this inherent noise necessitates a quantity of data orders of magnitude larger than similar studies of other species.

The fine timescale of our trajectory data (comprised of almost 20 000 individual locust trajectories, resolved at 25 frames per second) allowed the first analysis of individual locust motion in the field. We classified motion into three distinct modes: stationary, walking, and hopping. Intermittent motion was previously quantified in laboratory studies of different locust species (the desert locust) [15], and has typically been considered a binary between “stop” and “go”. Our evidence suggests a significant difference between the two moving states (walking and hopping) for the Australian plague locust. In theoretical models, intermittent motion provides a collective mechanism for density regulation [13, 17, 19]. By including hopping in future models, our findings may help to reproduce the dense fronts that occur in hopper bands of the Australian plague locust.

Our data has revealed novel anisotropies in the positions of neighbors around focal locusts in a natural setting. In particular, we find a lower likelihood of frontal neighbors around moving (walking and hopping) focal locusts. This represents a significant advance over the only other study that examined interactions of Australian plague locust within a hopper band [23]. Relying on manual data collection, that study computed a rotationally symmetric distribution from less than 20 000 neighbor positions in a disc of radius greater than 28 cm. Automated particle tracking enabled us to collect nearly 20 000 000 neighbor positions in a disc of radius 14 cm. After accounting for the difference in area, this represents an increase in number of data points by four orders of magnitude. In turn, this revealed the previously invisible anisotropies now clearly apparent in Figures 4.

These anisotropies have not been produced by models of locust interaction and hopper band collective motion. Previous models of locust interaction are either isotropic [17, 18] or based on the idea of escape-and-pursuit [22]. As shown explicitly by Buhl et al. [23], the escape-and-pursuit paradigm generates an anisotropy in neighbor density where localized high-density areas appear in front and behind the focal locust. This contrasts with the anisotropy found in our analysis. Models with isotropic interaction produce isotropic neighbor densities, also contrasting with our findings. New mechanisms of locust interaction are likely necessary to emulate our findings.

One possible explanation for our anisotropy results could be external factors such as prevailing winds or orientation of the sun. However, direct field observations have noted no correlation between a band’s direction and either of these [16, 24]. A second possible explanation could be that locusts themselves do not have an isotropic shape. Typically, locust bodies are much longer (head to end of abdomen) than they are wide. In a recent study of the desert locust, Gorbonos et al. [26] report that relative density distributions become isotropic after applying a correction to nearby neighbor data. In contrast, our anisotropy results for the Australian plague locust persist under similar corrections, see Appendix G for details. The simplest interpretation for these differing results is that the individual interactions differ between these two species of locust.

### Anisotropies in other animals

Most quantitative studies of animal groups reveal structure in the position of nearby neighbors. In contrast to our findings for locusts, higher densities are observed ahead and behind the focal individual in the Serengeti wildebeest [6], surface-swimming surf scoters [5], and some species of fish [11]. This type of anisotropy is typically associated with following behavior. Various species of fish display a range of likely relative neighbor positions, including at a diagonal or with no preference [27] and laterally [3, 28]. Starlings keep their nearest neighbors on their left and right, but this preference fades when considering all neighbors in a given radius [10].

We found an area of lower density ahead of moving focal locusts, see Figure 4 (center, right), which is a unique feature when compared with other species. Quantifying this in the front-back asymmetry, Figure 5 (dashed blue curve), we established its presence at distances less than 7 cm around walking locusts and at distances less than 2 cm and greater than 7 cm for hopping locusts. Simultaneously, we found that distributions of neighbors around moving locusts exhibit a four-fold anisotropy similar to that of starlings, with lateral areas of high density, see Figures 4 (right), and 5 (solid orange curve). Various explanations have been given for the higher occurrence of lateral neighbors for airborne birds and some fish. Ballerini et al. [4] suggests that this four-fold anisotropy could be due to a hydro/aerodynamic advantage, anisotropic vision, or a mechanism to avoid collisions in a high-speed group. Since these juvenile locusts do not fly, they would derive no significant aerodynamic benefit. Since locusts have essentially 360^◦^ vision [29], limited sight lines are not a likely explanation. Given the additional presence of front-back asymmetry around moving locusts, we suggest that the mechanism causing the anisotropies revealed here is one of collision avoidance.

Collision avoidance has been observed in other species exhibiting collective behavior. Above we noted it as a potential mechanism explaining the relative position of lateral neighbors in starlings [4]. In surf scoters [5], where high-density areas were found directly in front and behind focal individuals, examining the deviation from the mean velocity in the presence of frontal neighbors revealed that these ducks tend to follow frontal neighbors with a preferred following distance, slowing down to avoid collisions. Similarly, Katz et al. [11] found that golden shiners modulate their speed to avoid collisions while following neighboring fish. These following behaviors are among the best-documented examples of collision avoidance in collective behavior that we are aware of, but are qualitatively different from the case of locusts. Locusts appear to have far less control over their speed, likely due to the incredibly high strength to mass ratio characteristic of insects. We hypothesize that they instead tend to change their direction or type of motion to avoid collisions.

### Collision avoidance in locusts

Collision avoidance behaviors are well-documented through laboratory experiments in adult locusts; see a review by Fotowat and Gabbiani [30]. Both neural and behavioral responses occur from visual looming stimuli [31, 32] and have been particularly associated with collision trajectories [33, 34]. Stimuli in these studies had an area of at least 5 cm × 5 cm and responses were associated to a particular size of retinal image on the eye of the locust [31]. Response behaviors have included avoidant gliding for tethered flying locusts [35] and jumping away from an incoming stimulus for locusts on the ground [36]. This body of work establishes that adult locusts have the physiology necessary to sense nearby obstacles and react to avoid collisions. Our empirical findings suggest that juvenile locusts may exhibit similar visual sensing and motion adjustment; in our case in the midst of a naturally-occurring swarm, albeit while marching en mass on the ground rather than in flight. It is unclear whether this behavior may be due to the same neural circuits well-studied in adult locusts, opening up new behavioral and neurological questions.

### Recommendations for locust models of collective motion

Models of hopper band movement continue to advance towards the capability to predict the direction and distance that a given band will travel. The aim of such predictive models is to inform efficient control strategies for agricultural industry and government management agencies, possibly in real time. Predicting the likely trajectory and collective momentum of a threatening band can aid in conducting efficient surveys or implementing direct control strategies, such as pesticide barrier spraying [37]. Our results reveal state-dependent elements of locust motion and interaction during marching that are yet untested by current models and provide promising explanations for collective structure.

To start, we suggest a three-state model for capturing locust motion. The high level of accuracy (estimated at 85.0 %) of our method for classifying motion states supports this framework. Moreover, each motion state comprised a significant fraction (nearly 25%) of the data derived from locusts naturally marching in the field. This makes it difficult to justify omitting any one motion state from a realistic model. Additional evidence comes from the clean division of our distribution of individual speeds shown in Figure 3. A possible implementation could use a discrete time Markov process to dictate switches between stationary, walking, and hopping states. Going further, our distributions of speeds for each state could add biologically-realistic individual variation to such a model.

Secondly, we suggest modeling locust interactions with mechanisms for collision avoidance. By contrast, existing models of locust interaction either treat equidistant neighbors the same (i.e. are isotropic) or implement the escape-and-pursuit paradigm [14, 22] (where motion is driven by chasing behavior). As noted above, we find that moving locusts (both walking and hopping) have a high-density of neighbors on the left and right. A first possible mechanism for this anisotropy might be to implement a preference for lateral neighbors. However, this alone would likely not explain the additional front-back anisotropy we observe around moving locusts.

The area of lower neighbor density immediately in front of walking locusts (Figure 4, center) suggests that when walking locusts see another individual in front of them, they react to avoid a direct collision. This behavior could appear in a model by using some anisotropic kernel function when updating heading direction according to nearby neighbors and a lengthscale could be drawn from Figure 5 (center).

The anisotropy around hopping locusts (Figure 4, right) suggests a slightly different mechanism. Indeed, juvenile hoppers cannot change their motion mid-jump. Noting that the low-density area ahead of hopping locusts is narrower and more elongated than ahead of walking locusts, we suggest that one possibility is that locusts visually inspect a long and narrow area ahead of them before hopping and are more likely to hop when the path ahead is clear. Supporting this suggestion, there is a slight increase in front-back anisotropy for distances greater than 7 cm ahead of hopping locusts, see Figure 5 (right). This lengthscale also provides modelers with a valuable parameter when implementing such a mechanism, which could be encoded into a front-neighbor-dependent probability to switch out of the hopping state.

Recent modeling studies by Taylor et al. [38] and Krongauz and Lazebnik [39] have specifically noted the impact collision avoidance interactions can have on collective behavior. We hope that future models of collective motion in locusts will incorporate some of these specifics in order to determine their effect on the collective structure and function of the hopper band. Such predictions at the band level can then be tested in future field studies.

### Further Refinements

There are two limitations of our study that bear acknowledgement and provide opportunity for further investigation. The first is that our particle tracking implementation only identifies locust position. Consequently, we must infer heading direction from velocity. Since locusts move forward, almost never backwards or sideways, we are confident in the assumption that body orientation is equivalent to heading direction, i.e. direction of motion. The remaining issue is that stationary locusts do not have a well-defined heading direction. Via smoothing and interpolation we assigned a heading direction where we could confidently do so, but applying particle tracking software that collects body orientation, such as the newest release of TrackMate [40], may be preferable.

Secondly, we employed a supervised classification method for determining motion state. While we achieved a high degree of accuracy, a classification that makes use of unsupervised learning might uncover additional motion states that are not immediately apparent to the human eye while watching the recorded footage. For instance, only after watching footage in detail did we begin to notice non-moving locusts rotating their body’s orientation without advancing in any direction. This often occurred after a locust made an especially large jump. Particularly for small hoppers, these large jumps ended with a crash landing so that the insect’s orientation was no longer aligned with the prevailing direction of the band’s motion. The locust would rotate to align itself with passing neighbors before hopping again. Especially in conjunction with tracking software that records body orientation, identifying this behavior as distinct from other hopping might provide even stronger evidence of collision avoidance or novel patterns of interaction.

### Towards predictive modeling

In addition to informing theoretical models of collective behaviour, our empirical investigation of individual mechanisms contributes to the development of a well parametrized predictive locust movement model. With the addition of our findings, we are ever closer to predicting collective band behavior from evidence-based mechanisms for individual motion and interaction.

## 4 Methods

### 4.1 Recording hopper bands

We recorded video footage of eight distinct hopper bands of Australian plague locust (APL), *Chortoicetes terminifera*, during 3–10 of November 2010 near Hillston, New South Wales, Australia. The Australian Plague Locust Commission directed us to the area and put us in contact with the local control agency in Hillston who then took us to potential study sites. Site locations were S33.54733 E145.06678 (for bands 1–3) and S33.20745 E145.09763 (band 4–8). The hopper bands were composed of late-instar juveniles (3rd to 5th). Of the eight bands recorded, we analyzed four for this study.

Our recording procedure was similar to that described by Buhl et al. [12]. We mounted a camera on a tripod so that it pointed vertically downwards, with a view angle approximately perpendicular to the ground. We extended the tripod’s central column so that its legs did not obstruct the field of view. This resulted in a recorded area of the ground approximately 0.6 m^2^ using a Panasonic camcorder which recorded in 1080i.

For most recordings, we placed the tripod in the center of a marching band. Placing the tripod often caused a temporary disturbance, which we allowed to dissipate before beginning recording once the natural flow of marching had resumed. In one case, for band 1, when the location of the hopper band was known and accessible, we set up the tripod ahead of the band allowing for a full recording of the center of the band from front to back. For consistency, in this case we analyzed footage from after the front had passed the camera.

For each band, we chose the recording area to be flat and devoid of vegetation. We used areas located away from major obstacles that could prevent or impede locust marching such as trees, creeks, and patches of dense vegetation. We placed a sheet of plywood in the camera’s view frame to provide a uniform background. We recorded the scale by placing a ruler in the field of view at the beginning of the video, or else by the known dimensions of the plywood (120 cm × 60 cm).

For the purposes of this study, we selected four recordings where we observe sustained marching. Each video has a resolution of 1920 × 1080 pixels and consists of 25 interlaced frames per second. For this study, a total of twenty-seven minutes of footage was analyzed, representing 24 300 frames. For a sample of our video footage, see Appendix A.

### 4.2 Extracting numerical trajectories (via motion tracking with Track-Mate)

We analyzed the footage using particle-tracking software TrackMate [25], a plugin for ImageJ. This software includes a suite of established particle detection and trajectory-linking algorithms, along with a user-friendly GUI and interoperability with Matlab for scripted batch tracking. For more on particle-tracking software and to see how an early version of TrackMate performed, see [41]. For full details on our video processing, tracking algorithms and parameters, and how we evaluated TrackMate’s accuracy, see Appendix C.

### 4.3 Inferring motion

From the trajectory data for each locust, we inferred instantaneous velocity and decomposed it into speed and heading direction. To accurately compute these quantities we first cleaned the trajectory data including a spatial transformation to correct for the camera angle and smoothing the position data. For more details on data cleaning, see Appendix D.

After smoothing the position data, we computed velocities using a central difference method. Speed was calculated as the magnitude of the velocity. Taking the angle of the velocity in a standard coordinate system (*x*-axis to the right, *y*-axis up) we calculated heading direction. Note that heading direction is not well-defined for an unmoving locust. In fact, small fluctuations in position (inherited from automatic tracking) can produce large fluctuations in the heading direction. To account for this, we used linear interpolation to recompute the heading direction for stationary locusts after we classified their motion states.

### 4.4 Measuring collective marching

Since we wish to analyze the behavior of individuals during marching, we compute three quantities that are associated with this collective behavior. In each frame of video, we compute the density *D* by counting the number of locusts detected and dividing by the physical area in the frame. Typically, a density greater than 20 locusts*/*m^2^ has been associated with marching [42, 13]. We also compute the polarization *P* as the length of the average of the direction vectors (cos *φ_i_*, sin *φ_i_)*, where *φ_i_* is the heading direction of the *i*^th^ locust. Polarization is a commonly-used order parameter ranging from 0 (completely disordered) to 1 (completely aligned). Marching locusts have demonstrated polarization values between 0.6 and 0.9 [12]. To supplement polarization, we also compute an index that captures an adaptation of Boltzmann entropy. This entropy index *E* varies from 0 (completely aligned) to 1 (completely disordered) and was originally described by Baldassarre [43]. For marching locusts, the same entropy index has been reported between 0.75 and 0.9 [12] and we follow the same implementation therein. For details on formulas or computations of density, polarization, or entropy index refer to Appendix F.

### 4.5 Classifying motion state

Watching the raw footage, we observed locusts moving in one of three distinct motion states:

1. Stationary – locusts do not advance in any direction, may rotate their body’s orientation
2. Walking – locusts advance relatively slowly without leaving the ground, with short pauses separated by ∼ 1 s of motion
3. Hopping – locusts advance quickly through erratic jumps, sometimes with pauses of up to ∼ 1 s between jumps.

Distinct motion states have been previously recognized in locusts, including all three of these [24, 1, 15]. We manually classified all locusts in the two manually-tracked 10 s clips, creating our training and test data for automatic classification.

We implemented a support vector machine (SVM) to automatically classify each locust in each frame into one of the three motion states. SVMs are popular classification tools from machine learning – see the book by James et al. [44, Ch. 9] for an accessible introduction. SVMs compute an optimal boundary between each class of data points in a training data set, then use those boundaries to classify the remaining data. The boundaries depend only on the data points closest to them, called the *support vectors*, so the method is not sensitive to variations in well classified data points. For our application, we used a nonlinear boundary via a Gaussian kernel function. Computing the boundary amounts to an optimization problem where misclassified data points are assigned a penalty value based on their distance from the boundary in a higher-dimensional space. We implemented our SVM using Matlab’s `fitcecoc()` function.

After initial exploration of the data, we chose four trajectory features to classify locust motion states in each frame. For each locust in each frame of video, we centered a time window on the current frame and computed the following summary statistics for classification:

1. The instantaneous speed (after the smoothing described in) distinguishes locusts that are stationary at the current time;
2. the magnitude of average velocity distinguishes a stationary locust from one that has paused between jumps;
3. the standard deviation of speed distinguishes between hopping (fast and intermittent speed) and walking (slow and almost constant speed); and
4. the minimum of the forward and backward maximum speeds helps to discern between stationary and hopping locusts near the beginning or end of their trajectories in in these states.

We computed the latter three using a moving time window of 15 frames, equivalent to 0.6 s. The time window was chosen by tuning the classification accuracy of our SVM on the training data. To avoid overfitting, we optimized various internal hyperparameters of our SVM using cross-validation on the training data. We trained the SVM using the same data set and found that it correctly classified 86.8 % of the training data. We then evaluated the accuracy of our SVM on the second ground-truth data set and found correct classification for 85.0 % of the test data. Given the magnitude of the whole data set, this represents a level of accuracy that should easily distinguish trends from noise. See Appendix A for a sample video that illustrates the accuracy of our motion classification.

### 4.6 Quantifying anisotropy in neighbor density

Using the position and heading direction data, we aggregated relative positions of neighbors into density distributions using a methodology similar to Buhl et al. [23]. For each focal locust in a frame, we computed the relative position of each neighbor as its position in a coordinate frame with the focal individual at the origin, the *y*-axis pointing from tail to head, and the *x*-axis protruding to its right. We avoid biases introduced by the edges of the frame by taking two precautions. First, we only considered neighbors within 14 cm of the focal locust. This distance includes the only estimate we know of for the interaction range between locusts, which is 13.5 cm [23]. Second, we used the Hanisch correction [45, 46], which ignores any neighbor at a distance further from the focal individual than the nearest edge. We discretized this distribution of relative neighbor positions into square bins with side length Δ*x* = Δ*y* = 0.5 cm. We normalized the counts in each bin by the average density over all focal locusts divided by the average density in the whole area. The effect of this normalization factor is that as the distribution approaches homogeneity, the value in each bin approaches 1. We call these distributions *relative neighbor densities* and plot them as two-dimensional maps where color indicates density, see Figures 1 (right) and 4. This relative density can equivalently be thought of as the likelihood of finding a neighbor in a given position relative to a focal individual.

We also extracted the angle of each relative neighbor position and examined these as a distribution on the circle. An isotropic distribution of neighbors would correspond to a uniform distribution of their angles on the circle, so we used the Hodges-Ajne test for uniformity as described and implemented in the Matlab Circular Statistics Toolbox [47]. We quantify the non-uniformity of these distributions using *trigonometric moments* 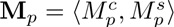 as described by Jammalamadaka and SenGupta [48]. In Figure 5, we examine how particularly relevant trigonometric moments vary by distance from the focal individual by looking at subsets of neighbors in concentric annuli. For details, see Appendix F.

## Supporting information

Sample Video

Sample Video with Data

## Acknowledgements

We thank the Australian Plague Locust Commission, the New South Wales Department of Primary Industries, Darron Cullen, Katie Robinson and Feng Hu for their help in the field. We thank Michael Culshaw-Maurer and Christopher Strickland for valuable conversations on how best to handle and analyze the large quanity of data. JW thanks Andrew Bernoff for support and guidance from the conception of the analysis to its conclusion. JL thanks Suzanne Amador Kane for her insight and mentorship throughout a preliminary version of the data investigation. JW was supported by an NSF Postdoctoral Fellowship Grant DMS–1902818. JL was supported by the HMC Data Science REU, NSF Grant DMS–1757952. AK, SL, and TS were supported by the Undergraduate Research Opportunities program at Harvey Mudd College. SJS, GAS and JB were funded by an ARC Linkage LP150100479. JB was funded by an ARC Future Fellowship FT110100082.

### Appendices A Sample video footage

Here we present links to a 10 s video clip from our raw video footage. We manually tracked the locusts in this clip to create the “ground-truth” test data that tuned the automatic tracking process and trained the motion classification machine.

1. Sample video footage showing juvenile Australian plague locusts, *Chortoicetes terminifera*, marching in a hopper band in the field. Recordings were captured by a camera mounted on a tripod with an extended arm and directed vertically down. A sheet of plywood was used to provide the uniform background. Locusts intermittently stop, walk, and hop as they advance in a common direction. • https://youtu.be/kTuzFzHl7Mo
2. Sample video with final processed data superimposed. Positions (circles) were obtained by automatic tracking. Heading directions (lines) were inferred from motion. Colors indicate motion state: stationary (red), walking (yellow), and hopping (green). Motion states were classified by a support vector machine operating on motion data from a local time window of each locusts trajectory. • https://youtu.be/ASOetHW2eVk

### B Tables

**Table 1:**
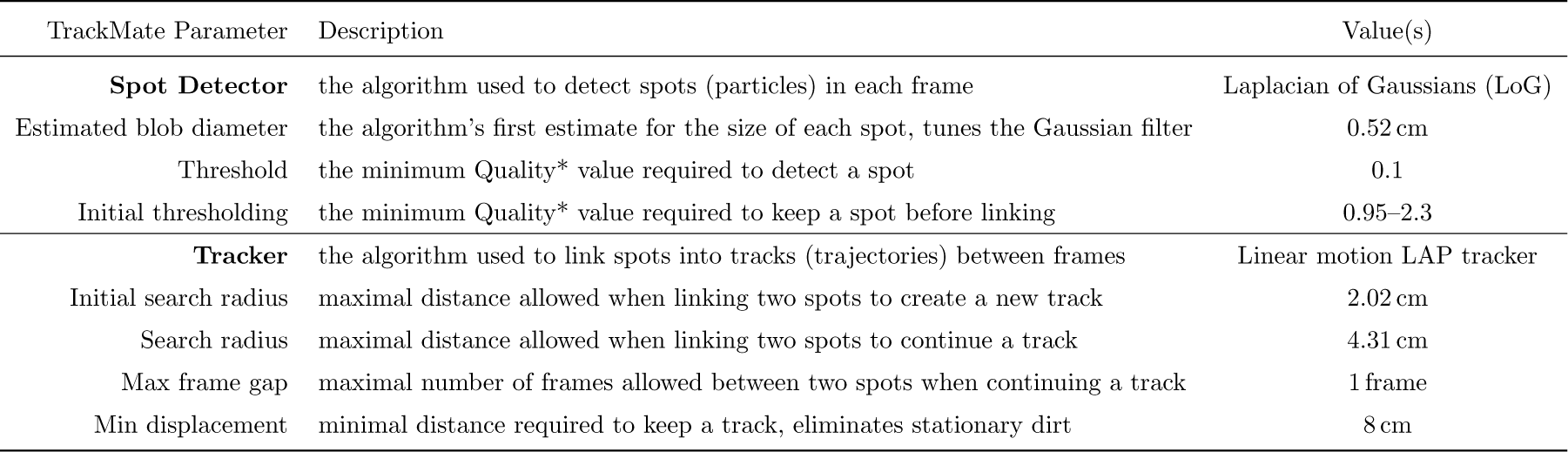
Tracking parameters for TrackMate, descriptions, and values used in our tracking. *Quality is defined as the local maximum value when the LoG filter is applied, see [25, Section 5.1.2] for details.

| TrackMate Parameter | Description | Value(s) |
| --- | --- | --- |
| **Spot Detector** | the algorithm used to detect spots (particles) in each frame | Laplacian of Gaussians (LoG) |
| Estimated blob diameter | the algorithm’s first estimate for the size of each spot, tunes the Gaussian filter | 0.52 cm |
| Threshold | the minimum Quality* value required to detect a spot | 0.1 |
| Initial thresholding | the minimum Quality* value required to keep a spot before linking | 0.95–2.3 |
| **Tracker** | the algorithm used to link spots into tracks (trajectories) between frames | Linear motion LAP tracker |
| Initial search radius | maximal distance allowed when linking two spots to create a new track | 2.02 cm |
| Search radius | maximal distance allowed when linking two spots to continue a track | 4.31 cm |
| Max frame gap | maximal number of frames allowed between two spots when continuing a track | 1 frame |
| Min displacement | minimal distance required to keep a track, eliminates stationary dirt | 8 cm |

**Table 2:** Indicators of collective marching for the full data set and divided by band. Typically, densities greater than 20 locusts*/*m^2^ are associated with marching. Marching locusts have demonstrated polarization values in the range 0.6–0.9 and entropy index values of 0.75–0.9 [12]. Values are given as mean plus/minus one standard deviation. Definitions of density, polarization, and the entropy index are given in Measuring collective marching and a subsample of the full distributions is plotted in Figure 2. Note that values for each band are consistent with marching behavior.

| | $N$<br>(locusts) | Density<br>(locusts/m<sup>2</sup>) | Polarization | Entropy Index |
| --- | --- | --- | --- | --- |
| all bands | 3369723 | 141.6 ± 71.0 | 0.82 ± 0.062 | 0.76 ± 0.058 |
| band 1 | 1135257 | 226.7 ± 65.2 | 0.84 ± 0.040 | 0.73 ± 0.033 |
| band 2 | 921094 | 145.9 ± 19.1 | 0.81 ± 0.043 | 0.77 ± 0.023 |
| band 3 | 979797 | 150.4 ± 35.7 | 0.80 ± 0.059 | 0.72 ± 0.038 |
| band 4 | 333575 | 55.5 ± 21.4 | 0.83 ± 0.082 | 0.82 ± 0.058 |

**Table 3:** Percent of data in each motion state and speeds divided by motion state for the complete data set and for each band. The percent of motion states varies significantly by band, probably due to environmental conditions such as temperature, but there is consistently a significant portion in each motion state. Speeds are given as mean plus/minus one standard deviation. Full distributions of speed are plotted as histograms in Figure 1 (left) and Figure 3.

| | $N$<br>(speeds) | Motion State (percent) | | | Speed, mean ± SD (cm s<sup>-1</sup>) | | | |
| --- | --- | --- | --- | --- | --- | --- | --- | --- |
|  |  | Stationary % | Walking % | Hopping % | All | Stationary | Walking | Hopping |
| all bands | 3332137 | 26.0 | 24.7 | 49.2 | 6.5 ± 8.4 | 0.2 ± 0.7 | 2.7 ± 2.2 | 11.8 ± 9.3 |
| band 1 | 1119569 | 12.2 | 26.2 | 61.6 | 8.4 ± 8.8 | 0.2 ± 0.8 | 3.1 ± 2.3 | 12.4 ± 9.1 |
| band 2 | 914361 | 39.9 | 28.1 | 32.0 | 4.1 ± 6.5 | 0.1 ± 0.5 | 2.6 ± 2.0 | 10.4 ± 8.1 |
| band 3 | 968173 | 28.8 | 18.5 | 52.7 | 6.3 ± 8.9 | 0.2 ± 0.9 | 2.1 ± 2.1 | 11.1 ± 10.0 |
| band 4 | 330034 | 26.6 | 28.7 | 44.7 | 7.4 ± 8.8 | 0.1 ± 0.4 | 3.2 ± 2.3 | 14.4 ± 8.9 |

**Table 4:** Trigonometric moments computed from distributions of angles of relative neighbor positions within 7 cm of focal locusts in each motion state, see Quantifying anisotropy in neighbor density for full definitions. The number of separated concentrations is given by *p*. Overall anisotropy is characterized by |**M**_p_| with direction given by *φ_p_* (and *φ_p_* + 2*π/p*). Anisotropy aligned with the focal locust’s heading direction is given 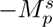 and perpendicular to the focal’s heading is given by 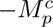. Thus *front-back asymmetry* is measured by 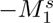 and *four-fold anisotropy* perpendicular to the direction of motion is measured by 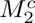. Bolded values correspond to features we plot against the distance from the focal locust in Figure 5. Overall anisotropy |**M**_p_| is significantly lower around stationary locusts than either moving state (for both *p* values). The angle *φ*_1_ is approximately −π/2 around walking locusts, indicating that the primary two-fold anisotropy is front-back. The angle *φ*_2_ is approximately 0 around both walking and hopping locusts, indicating that the primary four-fold anisotropy is perpendicular to the heading direction of the focal individual.

| All bands | $N$<br>(angles) | Two-fold anisotropy, $p = 1$ | | | | Four-fold anisotropy, $p = 2$ | | | |
| --- | --- | --- | --- | --- | --- | --- | --- | --- | --- |
| | | $\vert \mathbf{M}_1\vert $ | $\phi_1$ | $-M_1^s$ | $M_1^c$ | $\vert \mathbf{M}_2\vert $ | $\phi_2$ | $-M_2^s$ | $M_2^c$ |
| All Motion States | 6572930 | 0.0119 | -0.5341 | 0.0061 | 0.0102 | 0.0282 | -0.0060 | 0.0002 | 0.0282 |
| Stationary | 1505672 | 0.0040 | -2.5370 | **0.0023** | -0.0033 | 0.0146 | -0.2191 | 0.0032 | **0.0142** |
| Walking | 1527233 | 0.0173 | -1.5242 | **0.0173** | 0.0008 | 0.0248 | -0.0670 | 0.0017 | **0.0247** |
| Hopping | 3536882 | 0.0203 | -0.1393 | **0.0028** | 0.0201 | 0.0358 | 0.0492 | -0.0018 | **0.0357** |

### C TrackMate Details

#### Video preprocessing

Before tracking, we preprocessed the videos using ffmpeg [49] and Fiji [50] (ImageJ). Using ffmpeg we clipped the videos into computationally manageable lengths of 1 min. These clips were deinterlaced and saved to a nv12 AVI file format, which can be imported into ImageJ. Next, we loaded these clips into Fiji, a distribution of ImageJ that ships with the TrackMate plugin. In Fiji, we further processed the clips by applying a grayscale filter, inverting the colors, subtracting the median image (which effectively removes the nonmoving background), adjusting the brightness and contrast, and blurring (which smooths over remaining dirt particles and makes locust images more circular). This processing procedure reduced background noise in the video and prepared it for TrackMate, which performs best on approximately circular bright particles against a dark background.

#### Tracking algorithms and parameters

A full list of the tracking parameters we used appears in (Table 1); here we discuss the two major algorithms. The first step in particle tracking is to examine each frame and detect the location of particles (called “spots” in TrackMate). We used TrackMate’s Laplacian of Gaussians for our spot detection algorithm because it performs best for spots with diameter 5–20 pixels, which includes our blurred locusts. Mathematically, a Gaussian filter with standard deviation tuned by the expected spot radius is applied to the image. This smoothed image is then processed with a discrete Laplacian operator that detects local maxima. After spots have been detected in each frame, they are linked into trajectories (called “tracks”). TrackMate’s Linear Motion Tracker links spots assuming a roughly constant velocity between frames, which fits our video well because a typical locust reaction time of 0.03 s [51] is close to the time between our frames (0.04 s). To link spots between a pair of frames, this algorithm combines a cost-based assignment problem developed by Jaqaman et al. [52] with predictions from a Kalman filter for each locust. More technical details can be found in the TrackMate documentation [53].

#### Evaluating accuracy

TrackMate was also used for manual tracking to create two 10-s data sets, which we treat as our ground truth. These data include a total of 542 trajectories composed of 68 308 locust positions. We used the first data set to tune various parameters and the second to evaluate the accuracy of our data. We henceforth refer to the first as our *training data* and to the second as our *test data*.

To compare a ground-truth data set to its analogous data obtained from automatic tracking, we use similarity coefficients adapted from the Jacard Similarity Coefficient used in a challenge from the International Symposium on Biomedical Imaging [41]. For spot similarity we compute the Spot Similarity Coefficient

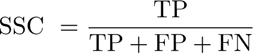

where TP is the number of true positive detections, FP is the number of false positives, and FN is the number of false negatives. Each of these is computed by an assignment problem between the two corresponding frames of each data set and a maximal assignment distance of 0.75 cm, less than 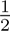 of a locust body length. The Link Similarity Coefficient is computed in a similar manner where we take pairs of frames from each data set. Each of these values ranges between 0 (no true positives) to 1 (perfect tracking). By tuning our tracking parameters we optimized the accuracy, achieving SSC = 0.97 and LSC = 0.96 for the training data. Evaluating these parameters against our test data we measured a spot accuracy of SSC = 0.91 and link accuracy of LSC = 0.90. This represents accurate tracking in a preponderance of the data. Given that we analyze 3 369 723 data points, this should be more than sufficient to distinguish signal from inherent noise. See Appendix A for a sample video illustrating the accuracy of our tracking.

### D Data

#### Data Cleaning

In each recording an attempt was made to place the camera aimed approximately perpendicular to the ground. However, we can observe that the angle was not exactly perpendicular by noting that the rectangular piece of plywood appears as an isosceles trapezoid in the video footage. For each recording, we constructed a spatial transformation that mapped the corners of the visual trapezoid (measured in pixels) to the corners of the actual dimensions of the plywood (120 cm × 60 cm). We applied this transformation to the position data for all trajectories of locusts marching in the corresponding recording. Next, we smoothed these trajectories through time.

Having such small mass, locusts can exhibit extremely large accelerations and we observed that locusts often changed their motion between frames of our video, which are separated by 0.04 s. Thus, more accurate velocity values can be obtained by smoothing the position data [54]. In order to choose an optimal interval length for the smoothing filter, we measured the timescale of significant autocorrelation in a manner similar to Ying et al. [55]. For a given trajectory, we computed the time series of distances traveled between frames and tested the null hypothesis that a unit root is present with an Augmented Dickey-Fuller (ADF) test. Of our ground-truth data, we found that 93.3 % of time series successfully rejected the null hypothesis and are therefore statistically stationary. We next computed the autocorrelation function with up to 20 time lags for each time in the trajectory. We judged a given time lag not to be significantly correlated with the original time if its autocorrelation function was less than two standard deviations from 0.

We recorded the time lag at the beginning of the first period of insignificantly correlated time lags. This represents the timescale after which a locust’s trajectory is not significantly correlated with itself. Across our ground truth data, we found a median correlation time of 3 frames (SD 4.8), or 0.12 s (SD 0.19 s). We therefore selected a weighted Gaussian filter over 8 frames as our smoothing function, which effectively links a given position only to four frames, or 0.16 s, in the future and four frames in the past.

#### Data Availability

Our final data set is available at the Dryad repository [56]:

- https://doi.org/10.5061/dryad.n02v6wwzz

The numerical trajectory data was extracted from video, cleaned, processed, and saved in Matlab’s .mat format. The repository also contains two subdirectories. The first has example material where one can execute our full data extraction, cleaning, and processing pipeline. This material includes the starting preprocessed video, intermediate processed video and .xml data files, and final example data. The second subdirectory contains the two ground-truth data sets obtained by manual tracking in TrackMate. The original video files are also included so that the manually-tracked .xml files can be opened in Fiji>Plugins>Tracking>Load a TrackMate File.

### E Code

The code used to extract, clean, and process the trajectory data, to conduct the analysis, and to generate the figures and tables is available on the public GitHub repository:

- https://github.com/weinburd/locust_trajectory_data

This repository has three levels of functionality.

1. By downloading only this repository, one can immediately run scripts to generate the figures and tables appearing in this paper from processed data stored in the repository itself.
2. By downloading this repository and the full data set in Appendix D, one can recreate the figures and tables directly from the final data set used in our analysis. Additionally, this option enables the user to implement our data cleaning and processing pipeline on a pair of example data sets and evaluate the tracking accuracy against our manually tracked ground-truth data.
3. By downloading this repository, the full data set, and installing Fiji locally on the users own computer, one can execute the examples described in (2) including the tracking process to extract numerical data from example video files. For this option, the user will need to ensure that their Fiji installation includes the TrackMate v6.0.3 plugin and enable the Matlab-ImageJ update site.

### F Mathematical Details

#### Density, Polarization, and Entropy

In a video frame containing *N* locusts, note that our measure of density

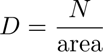

is relatively coarse. In our data, even the most highly populated frames do not approach the maximal packing density, so a more nuanced density measure would be computed locally in space. Still, our coarse measure is consistent with previous measurements.

We compute the polarization *P* as the length of the average of the direction vectors (cos *φ_i_*, sin *φ_i_)*, where *φ_i_* is the heading direction of the *i*^th^ locust. This quantity is

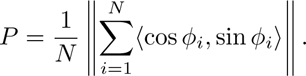

Polarization is a commonly-used order parameter ranging from 0 (completely disordered) to 1 (completely aligned). Marching locusts have demonstrated polarization values between 0.6 and 0.9 [12]. Polarization is known to be an ambiguous measure of alignment, since low values can result from either disorder or from two groups aligned in opposite directions. To supplement polarization, we also compute an index that captures an adaptation of Boltzmann entropy.

This entropy index *E* varies from 0 (completely aligned) to 1 (completely disordered) and was originally described by Baldassarre [43]. For marching locusts, this entropy index has been reported between 0.75 and 0.9 [12] and we follow the same implementation therein. For a frame with *N* locusts, we distribute their heading directions into an arbitrary number *C* = 72 of bins with width 2π/C. Letting *N_i_*be the number of angles in the *i*^th^ bin, we compute the entropy index as

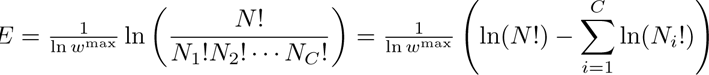

where *w*^max^ is the maximal value of 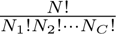 among all possible distributions of *N* heading directions into *C* classes, i.e. a nearly uniform distribution where each *N_i_* is either 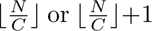. For *N >* 170, we use Stirling’s approximation ln(*N* !) ≈ *N* ln *N − N* + ^1^ ln(2π*N*) to avoid computational issues with large factorials. We acknowledge that the number of bins *C* can have a significant effect on the index. For a fixed *C*, however, the index *E* provides a stable relative measure of order. We use the same number of bins as Buhl et al. [12] because our goal is a direct comparison that establishes ordered marching for the locusts in our study.

#### Trigonometric Moments

We quantify the non-uniformity of the distribution of relative neighbor position angles using *trigonometric moments* as described by Jammalamadaka and SenGupta [48]. Given a set of *N* angles *φ_i_*, the *p*^th^ trigonometric moments are

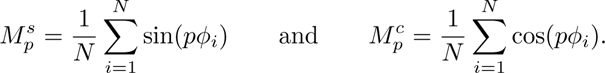

Mathematically, these are discretizations of the coefficients in the Fourier series expansion of a density function on the circle. In this way, they capture how much the data is concentrated into *p* groups, i.e. the Fourier modes on the circle. Physically, the vector 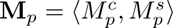 has magnitude 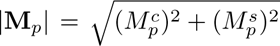, which provides a dimensionless measure that quantifies how much the data is concentrated into *p* equally-spaced groups. The vector **M***_p_* points in the direction *pφ_p_* where *φ_p_* is the angle of one of the *p* concentrations of data. A uniform distribution on the circle has |**M**_p_| = 0 and a completely concentrated distribution where all *φ_i_*’s are equal has |**M**_p_| = 1.

We are particularly interested in 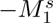 and 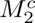. Recall the coordinate system in our relative neighbor density plots, see Figure 1 (right) and Figure 4. For the focal locust in the center and facing upwards, 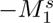 quantifies the *front-back asymmetry*. With the negative sign, this quantity is larger when there is a lower density in front and a higher density behind the focal locust. It is relatively stable to fluctuations in density on the left and right. By contrast, 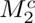 quantifies the *four-fold anisotropy* around the focal locust. It is largest when there are high-density concentrations to the left and right and lower-density areas in front and behind. In Figure 5, we examine how these quantities vary by distance from the focal individual by looking at subsets of neighbors in concentric annuli.

### G Anisotropy After Accounting for Locust Body Shape

A recent report studies neighbor density distributions in desert locusts [26] in a manner similar to ours. Findings therein complement our results with a similar anisotropic distribution including higher densities of neighbors on left and right of each focal individual. However, Gorbonos et al. [26] also find that this anisotropy vanishes after transforming their neighbor position data to account for the body shape of locusts. They suggest that a reasonable explanation for anisotropy in neighbor location among randomly interacting particles could be found by examining the shape of the particles. Locusts are not circular, but tend to be longer (from head to end of abdomen) than they are wide. Here we investigate this effect in our own data by examining two severe transformations of the neighbor data, each designed to compensate for the body shape of locusts. We demonstrate that the observed anisotropy in our results persists after both of these corrections.

We suppose that an average juvenile has a body length of 1.5 cm and a body width of 0.5 cm (adult Australian plague locust bodies generally range from 3 cm to 4.5 cm [57]). We consider two transformations to the relative position data of neighbors near each focal locust:

- rescale is given by the vector function 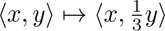; and
- extreme is given by the vector function *(x, y) 1→ (x, d)*, where *d* is the minimum forward distance between the focal and neighbor locust.

The first correction rescale is derived from approximating a locust’s body by an ellipse and then finding the transformation that changes that ellipse into a circle. The second correction extreme represents the most extreme assumption on locust body-shape: that each locust is a vertical line with no width. The corresponding transformation captures our interest in collision avoidance. In other words, we measure the minimum distance to collision if locust bodies were non-circular to an extreme.

The resulting relative neighbor density plots appear in Figure 6. These can be compared to Figure 4 but differ because they show the density only of the single neighbor nearest the focal locust. The top row includes no correction, note the anisotropies similar to those observed in Figure 4. The middle row applies the transformation rescale. Note that areas of higher density can still be observed to the left and right of walking and hopping focal locusts focal locusts. The effect is less visible but still present. Additionally, an area of lower density still appears clearly in front of walking locusts. In the bottom row we apply the transformation extreme and obtain predictably extreme results. Vertical distances have been subtracted, with a minimum of zero, from many nearest neighbors. This creates the lines on left and right of extremely high relative density – increasing the anisotropy of the neighbor distribution. Note that to capture details in the neighbors not exactly to left and right in these plots we adjusted the coloring to be on a logarithmic scale.

**Figure 6:**
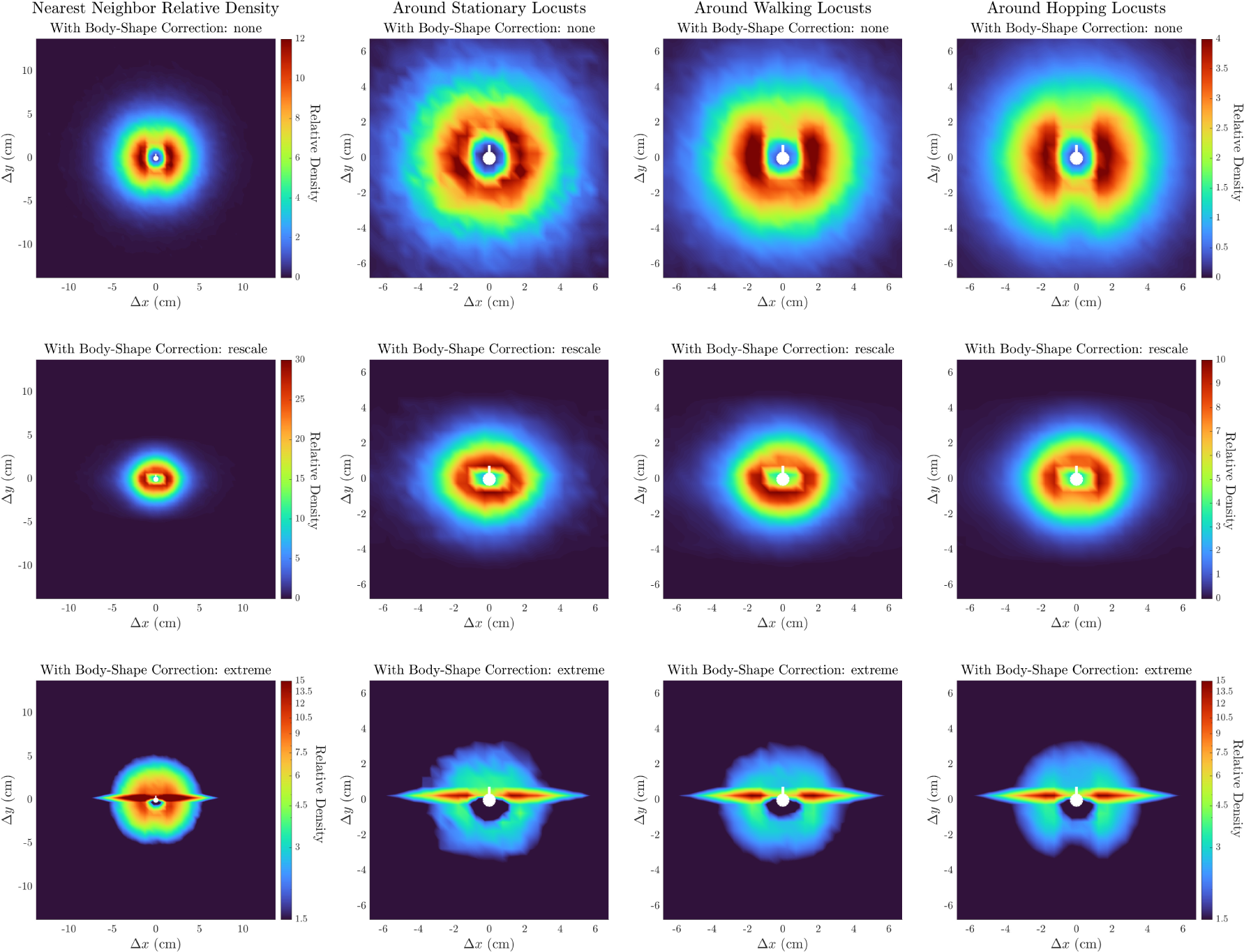
Relative neighbor density around focal locusts that are in (from left to right) any motion state, stopped, walking, and hopping. Plots are computed from *only single nearest neighbor* data with no correction (top row) and after transformations rescale (middle row) and extreme (bottom row). Focal locust (white marker), high (red) and low (blue) densities are as in Figure 4. Note that the anisotropy of the original neighbor densities are augmented but persist under both corrections for body shape. In particular, there are areas of higher density to left and right and an area of lower density in front of the focal locusts that are walking and hopping.

Acknowledging that the anisotropy is more subtle after applying the rescale transformation, we quantify the anisotropy before and after the correction in Figure 7. In order to facilitate comparison with concurrent work we here compute the proportion of nearest neighbors on the sides of each focal locust as described by Gorbonos et al. [26, Appendix D]. This measure is mathematically similar to the trigonometric moment 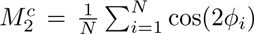, described more fully in Appendix F. They are related by replacing the continuous weight function cos(2φ) with the discontinuous indicator function

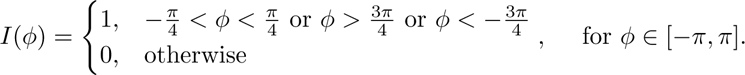

**Figure 7:**
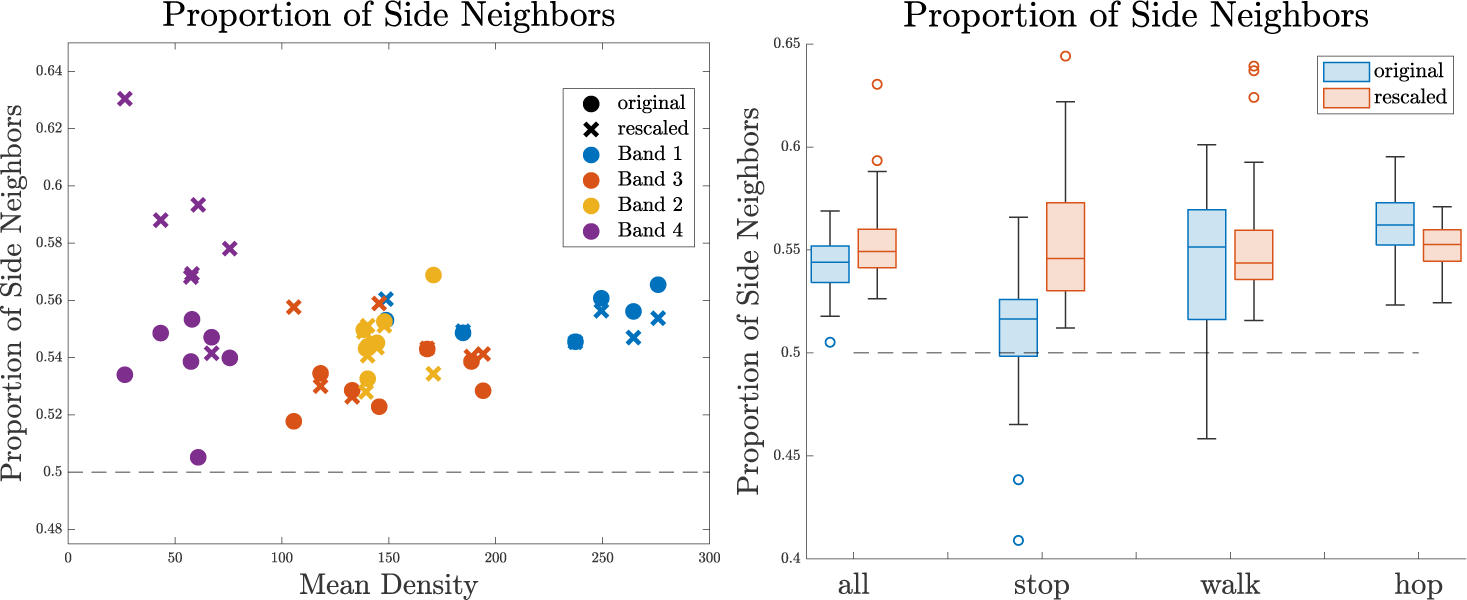
Proportion of nearest neighbors on the sides of focal locusts in each video by mean density (left) and for all videos by motion state (right). Box plots display median (center line), twenty-fifth and seventy-fifth quartiles (box bounds), remainder of the data (black whiskers), and outliers (circles). Note that the rescale transformation does not affect the proportion of side neighbors in a consistent fashion across all videos. All proportions of side neighbors after rescaling are bounded away from the fraction 0.5 corresponding to isotropy (dashed line).

With this substitution 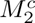 becomes the proportion of side neighbors 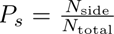. We consider only nearest neighbors within 4 cm of the focal locust because our data extends only 14 cm and we will contract these farthest distances by a factor of 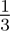.

The left-hand panel of Figure 7 shows this proportion *P_s_*for each video clip before and after the *rescale* transformation. No consistent increase or decrease in *P_s_* is discernible. The righ-hand panel shows a pair of box plots for focal locusts in each motion state. Each box plot captures the distribution of *P_s_*across all video clips. The dashed line at *P_s_*= 0.5 represents an isotropic distribution of neighbors. Note that when combining all motion states, the *rescale* transformation increases the overall anisotropy as measured by *P_s_*. For focal locusts that are walking and hopping, the distribution of *P_s_* shifts lower, but we still observe a significant gap between the whiskers and lower quartile and the dashed line *P_s_* = 0.5. In other words, the anisotropy decreases but persists.

## Notes

https://doi.org/10.5061/dryad.n02v6wwzz

https://github.com/weinburd/locust_trajectory_data

### Competing Interest Statement

The authors have declared no competing interest.

### Summary of Updates

Small changes throughout provide clarity, especially to highlight when additional detail can be found in an Appendix; A new panel was added to Figure 1 showing sample trajectories.

## References

[1] B. Uvarov. Grasshoppers and Locusts, volume 2. Cambridge University Press, London, UK, 1977.

[2] David Sumpter. Collective animal Behavior. Princeton University Press, 2010.

[3] Brian L. Partridge. The structure and function of fish schools. Scientific American, 246(6):114–123, 1982.

[4] Michele Ballerini, Nicola Cabibbo, Raphael Candelier, Andrea Cavagna, Evaristo Cisbani, Irene Giardina, Alberto Orlandi, Giorgio Parisi, Andrea Procaccini, Massimiliano Viale, and Vladimir Zdravkovic. Empirical investigation of starling flocks: a benchmark study in collective animal behaviour. Animal Behaviour, 76(1):201–215, jul 2008. doi: 10.1016/j.anbehav.2008.02.004. URL https://doi.org/10.1016%2Fj.anbehav.2008.02.004.

[5] Ryan Lukeman, Yue-Xian Li, and Leah Edelstein-Keshet. Inferring individual rules from collective behavior. Proceedings of the National Academy of Sciences, 107(28):12576–12580, 2010. ISSN 0027-8424. doi: 10.1073/pnas.1001763107. URL https://www.pnas.org/content/107/28/12576.

[6] Colin J. Torney, J. Grant C. Hopcraft, Thomas A. Morrison, Iain D. Couzin, and Simon A. Levin. From single steps to mass migration: the problem of scale in the movement ecology of the serengeti wildebeest. Phil. Trans. R. Soc. B, 373(1746):20170012, mar 2018. doi: 10.1098/rstb.2017.0012. URL https://doi.org/10.1098%2Frstb.2017.0012.

[7] Craig W. Reynolds. Flocks, herds, and schools: A distributed behavioral model. Computer Graphics (SIGGRAPH ‘87 Conference Proceedings), 21(4):25–34, 1987.

[8] T. Vicsek, A. Czirok, E. Ben Jacob, I. Cohen, and O. Shochet. Novel type of phase-transition in a system of self-driven particles. Phys. Rev. Lett., 75(6):1226–1229, August 1995.

[9] Felipe Cucker and Steve Smale. On the mathematics of emergence. Japanese Journal of Mathematics, 2(1):197–227, mar 2007. doi: 10.1007/s11537-007-0647-x. URL https://doi.org/10.1007%2Fs11537-007-0647-x.

[10] M. Ballerini, N. Cabibbo, R. Candelier, A. Cavagna, E. Cisbani, I. Giardina, V. Lecomte, A. Orlandi, G. Parisi, A. Procaccini, M. Viale, and V. Zdravkovic. Interaction ruling animal collective behavior depends on topological rather than metric distance: Evidence from a field study. Proceedings of the National Academy of Sciences, 105(4):1232–1237, 2008. ISSN 0027-8424. doi: 10.1073/pnas.0711437105. URL https://www.pnas.org/content/105/4/1232.

[11] Yael Katz, Kolbjørn Tunstrøm, Christos C. Ioannou, Cristían Huepe, and Iain D. Couzin. Inferring the structure and dynamics of interactions in schooling fish. Proceedings of the National Academy of Sciences, 108(46):18720–18725, 2011. ISSN 0027-8424. doi: 10.1073/pnas.1107583108. URL https://www.pnas.org/content/108/46/18720.

[12] Jerome Buhl, Gregory A Sword, Fiona J Clissold, and Stephen J Simpson. Group structure in locust migratory bands. Behavioral Ecology and Sociobiology, 65(2):265–273, 2011.

[13] J. Buhl, D.J.T. Sumpter, I. D. Couzin, J.J. Hale, E. Despland, E.R. Miller, and S. J. Simpson. From Disorder to Order in Marching Locusts. Science, 312:1402–1406, 2006. doi: 10.1126/science.1125142.

[14] Sepideh Bazazi, Jerome Buhl, Joseph J. Hale, Michael L. Anstey, Gregory A. Sword, Stephen J. Simpson, and Iain D. Couzin. Collective motion and cannibalism in locust migratory bands. Current Biology, 18(10):735–739, 2019/06/26 2008. doi: 10.1016/j.cub.2008.04.035. URL https://doi.org/10.1016/j.cub.2008.04.035.

[15] Gil Ariel, Yotam Ophir, Sagi Levi, Eshel Ben-Jacob, and Amir Ayali. Individual pause-and-go motion is instrumental to the formation and maintenance of swarms of marching locust nymphs. PLOS One, 9(7):e101636, 2014.

[16] L.R. Clark. Behaviour of Swarm Hoppers of the Australian Plague locust Chortoicetes terminifera (Walker). Commonwealth Scientific and Industrial Research Organization, Australia, 245:5–26, 1949.

[17] Jamila Dkhili, Uta Berger, Lalla Mina Idrissi Hassani, Säıd Ghaout, Ronny Peters, and Cyril Piou. Self-organized spatial structures of locust groups emerging from local interaction. Ecological Modelling, 361:26–40, October 2017. ISSN 03043800. doi: 10.1016/j.ecolmodel.2017.07.020.

[18] A. Bach. Exploring locust hopper bands emergent patterns using parallel computing. Master’s thesis, Université Paul Sabatier Toulouse III, June 2018.

[19] Andrew J. Bernoff, Michael Culshaw-Maurer, Rebecca A. Everett, Maryann E. Hohn, W. Christopher Strickland, and Jasper Weinburd. Agent-based and continuous models of hopper bands for the australian plague locust: How resource consumption mediates pulse formation and geometry. PLOS Computational Biology, 16(5):e1007820, may 2020. doi: 10.1371/journal.pcbi.1007820. URL https://doi.org/10.1371%2Fjournal.pcbi.1007820.

[20] Gil Ariel and Amir Ayali. Locust Collective Motion and Its Modeling. PLOS Computational Biology, pages 1–25, 2015. doi: 10.1371/journal.pcbi.1004522. URL https://doi.org/10.1371/journal.pcbi.1004522.

[21] Meir Paul Pener and Stephen J. Simpson. Locust Phase Polyphenism: An Update. In Advances in Insect Physiology, volume 36, pages 1–272. Elsevier, 2009. ISBN 978-0-12-374828-7. doi: 10.1016/S0065-2806(08)36001-9.

[22] Pawel Romanczuk, Iain D Couzin, and Lutz Schimansky-Geier. Collective Motion due to Individual Escape and Pursuit Response. Physical Review Letters, 010602:1–4, 2009. doi: 10.1103/PhysRevLett.102.010602.

[23] J Buhl, Gregory A Sword, and Stephen J Simpson. Using field data to test locust migratory band collective movement models. Interface Focus, 2:757–763, 2012.

[24] Peggy Ellis and Clifford Ashall. Field studies on diurnal behaviour, movement and aggregation in the desert locust (*schistocerca gregaria* forsk.). Anti-Locust Bulletin, 25:4–94, 1957.

[25] Jean-Yves Tinevez, Nick Perry, Johannes Schindelin, Genevieve M. Hoopes, Gregory D. Reynolds, Emmanuel Laplantine, Sebastian Y. Bednarek, Spencer L. Shorte, and Kevin W. Eliceiri. Track-mate: An open and extensible platform for single-particle tracking. Methods, 115:80–90, 2017. ISSN 1046-2023. doi: 10.1016/j.ymeth.2016.09.016. URL http://www.sciencedirect.com/science/article/pii/S1046202316303346. Image Processing for Biologists.

[26] Dan Gorbonos, Felix Oberhauser, Luke L. Costello, Yannick Günzel, Einat Couzin-Fuchs, Benjamin Koger, and Iain D. Couzin. An effective hydrodynamic description of marching locusts. 2023. URL https://arxiv.org/abs/2308.02589.

[27] Brian L. Partridge, Tony Pitcher, J. Michael Cullen, and John Wilson. The three-dimensional structure of fish schools. Behavioral Ecology and Sociobiology, 6(4):277–288, mar 1980. doi: 10.1007/bf00292770. URL https://doi.org/10.1007%2Fbf00292770.

[28] Daniel Grünbaum, Steven Viscido, and Julia K. Parrish. Extracting interactive control algorithms from group dynamics of schooling fish. In V. Kumar, N. Leonard, and A.S. Morse, editors, Cooperative Control. Lecture Notes in Control and Information Science, volume 309, pages 103–117. Springer Berlin Heidelberg, nov 2004. doi: 10.1007/978-3-540-31595-76. URL https://doi.org/10.1007%2F978-3-540-31595-7_6.

[29] Stephen M. Rogers, George W. J. Harston, Fleur Kilburn-Toppin, Thomas Matheson, Malcolm Burrows, Fabrizio Gabbiani, and Holger G. Krapp. Spatiotemporal receptive field properties of a looming-sensitive neuron in solitarious and gregarious phases of the desert locust. J. Neurophysiol, 103(2):779–792, feb 2010. doi: 10.1152/jn.00855.2009. URL https://doi.org/10.1152%2Fjn.00855.2009.

[30] Haleh Fotowat and Fabrizio Gabbiani. Collision detection as a model for sensory-motor integration. Annual Review of Neuroscience, 34(1):1–19, jul 2011. doi: 10.1146/annurev-neuro-061010-113632. URL https://doi.org/10.1146%2Fannurev-neuro-061010-113632.

[31] R. M. Robertson and A. G. Johnson. Retinal image size triggers obstacle avoidance in flying locusts. Naturwissenschaften, 80(4):176–178, apr 1993. doi: 10.1007/bf01226378. URL https://doi.org/10.1007%2Fbf01226378.

[32] John R. Gray, R. Meldrum Robertson, and Jessica K. Lee. Activity of descending contralateral movement detector neurons and collision avoidance behaviour in response to head-on visual stimuli in locusts. Journal of Comparative Physiology A, 187(2):115–129, apr 2001. doi: 10.1007/s003590100182. URL https://doi.org/10.1007%2Fs003590100182.

[33] S Judge and F Rind. The locust DCMD, a movement-detecting neurone tightly tuned to collision trajectories. Journal of Experimental Biology, 200(16):2209–2216, aug 1997. doi: 10.1242/jeb.200.16.2209. URL https://doi.org/10.1242%2Fjeb.200.16.2209.

[34] R. M. Robertson and A. G. Johnson. Collision avoidance of flying locusts: Steering torques and behavior. Journal of Experimental Biology, 183(1):35–60, oct 1993. doi: 10.1242/jeb.183.1.35. URL https://doi.org/10.1242%2Fjeb.183.1.35.

[35] Roger D. Santer, Peter J. Simmons, and F. Claire Rind. Gliding behaviour elicited by lateral looming stimuli in flying locusts. Journal of Comparative Physiology A, 191(1):61–73, nov 2004. doi: 10.1007/s00359-004-0572-x. URL https://doi.org/10.1007%2Fs00359-004-0572-x.

[36] Roger D. Santer, Yoshifumi Yamawaki, F. Claire Rind, and Peter J. Simmons. Motor activity and trajectory control during escape jumping in the locust locusta migratoria. Journal of Comparative Physiology A, 191(10):965–975, jul 2005. doi: 10.1007/s00359-005-0023-3. URL https://doi.org/10.1007%2Fs00359-005-0023-3.

[37] J. Holt and J.F. Cooper. A model to compare the suitability of locust hopper targets for control by insecticide barriers. Ecological Modelling, 195(3-4):273–280, jun 2006. doi: 10.1016/j.ecolmodel.2005.11.026. URL https://doi.org/10.1016%2Fj.ecolmodel.2005.11.026.

[38] Chris Taylor, Colin Luzzi, and Cameron Nowzari. On the effects of collision avoidance on emergent swarm behavior. In 2020 American Control Conference (ACC). IEEE, July 2020. doi: 10.23919/ acc45564.2020.9147834. URL https://doi.org/10.23919/ACC45564.2020.9147834.

[39] David L. Krongauz and Teddy Lazebnik. Collective evolution learning model for vision-based collective motion with collision avoidance. PLOS ONE, 18(5):e0270318, May 2023. ISSN 1932-6203. doi: 10.1371/journal.pone.0270318. URL https://doi.org/10.1371/journal.pone.0270318.

[40] Dmitry Ershov, Minh-Son Phan, Joanna W. Pylvänäinen, Stéphane U. Rigaud, Laure Le Blanc, Arthur Charles-Orszag, James R. W. Conway, Romain F. Laine, Nathan H. Roy, Daria Bonazzi, Guillaume Duménil, Guillaume Jacquemet, and Jean-Yves Tinevez. Bringing TrackMate into the era of machine-learning and deep-learning. bioRxiv 2021.09.03.458852, sep 2021. doi: 10.1101/2021.09.03.458852. URL https://doi.org/10.1101%2F2021.09.03.458852.

[41] Nicolas Chenouard, Ihor Smal, Fabrice de Chaumont, Martin Maška, Ivo F. Sbalzarini, Yuanhao Gong, Janick Cardinale, Craig Carthel, Stefano Coraluppi, Mark Winter, Andrew R. Cohen, William J. Godinez, Karl Rohr, Yannis Kalaidzidis, Liang Liang, James Duncan, Hongying Shen, Yingke Xu, Klas E. G. Magnusson, Joakim Jaldén, Helen M. Blau, Perrine Paul-Gilloteaux, Philippe Roudot, Charles Kervrann, Fraņcois Waharte, Jean-Yves Tinevez, Spencer L. Shorte, Joost Willemse, Katherine Celler, Gilles P. van Wezel, Han-Wei Dan, Yuh-Show Tsai, Carlos Ortiz de Soĺorzano, Jean-Christophe Olivo-Marin, and Erik Meijering. Objective comparison of particle tracking methods. Nature Methods, 11(3):281–289, Mar 2014. ISSN 1548-7105. doi: 10.1038/nmeth.2808. URL https://doi.org/10.1038/nmeth.2808.

[42] P. M. Symmons and K. Cressman. Desert locust guidelines. Technical report, Food and Agriculture Organization of the United Nations, Rome, 2001.

[43] Gianluca Baldassarre. Self-organization as phase transition in decentralized groups of robots: A study based on boltzmann entropy. In Advanced Information and Knowledge Processing, pages 157–177. Springer London, 2013. doi: 10.1007/978-1-4471-5113-57. URL https://doi.org/10.1007%2F978-1-4471-5113-5_7.

[44] Gareth James, Daniela Witten, Trevor Hastie, and Robert Tibshirani. An introduction to statistical learning, volume 112. Springer, 2013.

[45] K.H. Hanisch. Some remarks on estimators of the distribution function of nearest neighbour distance in stationary spatial point processes. Series Statistics, 15(3):409–412, jan 1984. doi: 10.1080/02331888408801788. URL https://doi.org/10.1080%2F02331888408801788.

[46] Peter Diggle. Statistical analysis of spatial and spatio-temporal point patterns. CRC press, 2013.

[47] Philipp Berens. CircStat: A MATLAB Toolbox for circular statistics. Journal of Statistical Software, 31(10), 2009. doi: 10.18637/jss.v031.i10. URL https://doi.org/10.18637%2Fjss.v031.i10.

[48] S Rao Jammalamadaka and Ashis SenGupta. Topics in Circular Statistics. World Scientific, apr 2001. doi: 10.1142/4031. URL https://doi.org/10.1142%2F4031.

[49] FFmpeg Developers. ffmpeg. Open Source, Accessed May Accessed 2020. URL https://ffmpeg.org/ffmpeg.html#Authors.

[50] Johannes Schindelin, Ignacio Arganda-Carreras, Erwin Frise, Verena Kaynig, Mark Longair, Tobias Pietzsch, Stephan Preibisch, Curtis Rueden, Stephan Saalfeld, Benjamin Schmid, Jean-Yves Tinevez, Daniel James White, Volker Hartenstein, Kevin Eliceiri, Pavel Tomancak, and Albert Cardona. Fiji: an open-source platform for biological-image analysis. Nature Methods, 9(7):676–682, jun 2012. doi: 10.1038/nmeth.2019. URL https://doi.org/10.1038%2Fnmeth.2019.

[51] T. Friedel. The vibrational startle response of the desert locust schistocerca gregaria. Journal of Experimental Biology, 202(16):2151–2159, aug 1999. doi: 10.1242/jeb.202.16.2151. URL https://doi.org/10.1242%2Fjeb.202.16.2151.

[52] Khuloud Jaqaman, Dinah Loerke, Marcel Mettlen, Hirotaka Kuwata, Sergio Grinstein, Sandra L Schmid, and Gaudenz Danuser. Robust single-particle tracking in live-cell time-lapse sequences. Nature Methods, 5(8):695–702, jul 2008. doi: 10.1038/nmeth.1237. URL https://doi.org/10.1038%2Fnmeth.1237.

[53] Jean-Yves Tinevez. TrackMate Manual, Aug 2016. URL https://imagej.net/media/plugins/trackmate/trackmate-manual.pdf.

[54] Kenichi Shibata and Takashi Amemiya. How to decide window-sizes of smoothing methods: A goodness of fit criterion for smoothing oscillation data. IEICE Transactions on Electronics, E102.C(2): 143–146, feb 2019. doi: 10.1587/transele.2018oms0003. URL https://doi.org/10.1587%2Ftransele.2018oms0003.

[55] Wenxia Ying, Gabriel Huerta, Stanly Steinberg, and Martha Zúñiga. Time series analysis of particle tracking data for molecular motion on the cell membrane. Bulletin of Mathematical Biology, 71(8):1967–2024, aug 2009. doi: 10.1007/s11538-009-9434-6. URL https://doi.org/10.1007%2Fs11538-009-9434-6.

[56] Jasper Weinburd, Stephen J. Simpson, Gregory A. Sword, and Jerome Buhl. Trajectory data for locusts in a hopper band [dataset]. Dryad. URL DOI:10.5061/dryad.n02v6wwzz.

[57] Fisheries Australian Government Department of Agriculture and Forestry. URL https://www.agriculture.gov.au/biosecurity-trade/pests-diseases-weeds/locusts/about/id-guide/description_of_adults/1_australian_plague_locust_chortoicetes_terminifera.

